# To be or not to be - Decoding the Trabecular Meshwork Cell Identity

**DOI:** 10.1101/2024.04.26.591346

**Authors:** Alice Tian, Hasna Baidouri, Sangbae Kim, Jin Li, Xuesen Cheng, Yumei Li, Rui Chen, VijayKrishna Raghunathan

**Affiliations:** Human Genome Sequencing Center, Baylor College of Medicine, Houston, Texas 77030, USA; Department of Molecular and Human Genetics, Baylor College of Medicine, Houston, Texas 77030, USA; Department of Biochemistry and Molecular Biology, Baylor College of Medicine, Houston, Texas 77030, USA; University of Houston, College of Optomtery, Houston, TX, 77204, USA

## Abstract

The trabecular meshwork within the conventional outflow apparatus is critical in maintaining intraocular pressure homeostasis. *In vitro* studies employing primary cell cultures of the human trabecular meshwork (hTM) have conventionally served as surrogates for investigating the pathobiology of TM dysfunction. Despite its abundant use, translation of outcomes from *in vitro* studies to *ex vivo* and/or *in vivo* studies remains a challenge. Given the cell heterogeneity, performing single-cell RNA sequencing comparing primary hTM cell cultures to hTM tissue may provide important insights on cellular identity and translatability, as such an approach has not been reported before. In this study, we assembled a total of 14 primary hTM *in vitro* samples across passages 1-4, including 4 samples from individuals diagnosed with glaucoma. This dataset offers a comprehensive transcriptomic resource of primary hTM *in vitro* scRNA-seq data to study global changes in gene expression in comparison to cells in tissue *in situ*. We have performed extensive preprocessing and quality control, allowing the research community to access and utilize this public resource.

## Background & Summary

Primary open angle glaucoma (POAG) is a devastating ocular disorder resulting in irreversible vision loss. While the precise etiology and progression of POAG are complex, reducing intraocular pressure (IOP) is the only modifiable risk factor for managing visual field loss. The trabecular meshwork (TM) in the anterior segment of the eye is a major site of egress for aqueous humor^1^. Dysfunction of the TM, contributed by tissue resident cells and the extracellular matrix (ECM), is thought to result in increased resistance to aqueous drainage and ocular hypertension (OHT)^2,3^. Conventionally, the TM is thought to be anatomically heterogeneous and is comprised of 3 regions: the juxtacanalicular region (JCT) within the cribiform region, corneoscleral meshwork, and uveoscleral meshwork, of which the area around the juxtracanalicular region (JCT) of the meshwork and the inner wall cells of the Schlemm’s canal is considered the major site of resistance to outflow^4–8^. The primary intervention to reducing OHT is achieved through targeting either reduction of aqueous production (e.g. *α*-adrenergic agonists, *β* -blockers, carbonic anhydrase inhibitor), or via increasing outflow via the unconventional pathway (e.g. prostaglandins, miotic & cholinergic agents). More recently, Rho kinase inhibitors and nitric oxide donor drugs that target the conventional outflow apparatus have been approved for use.

Understanding the molecular pharmacology and mechanism of action of drugs has often relied on primary TM cells cultured *in vitro* prior to conduction of *ex vivo* or *in vivo* studies. However, translation of *in vitro* efficacy to pre-clinical and subsequently clinical outcomes remains a challenge. Differences in tissue anatomy and ocular structures, physiology, microenvironment, and species all contribute to the inability to replicate the complexities of an organism *in vitro*. The physiology and anatomy of the TM is complex with functional heterogeneity observed in the form of segmental regions of high, intermediate and low flow^9^. Segmental heterogeneity is associated with changes in extracellular matrix composition, biomechanical properties, and molecular signaling pathways in cells^10–14^. To further understand this complex tissue, single cell transcriptomic studies demonstrate heterogeneity in cell types within the conventional pathway with 12-to-19 distinct cell types identified with region-specific expression of candidate genes to define cellular identity^15,16^. In contrast, *in vitro* cell culture of primary human TM (hTM) cells originate with isolation of these cells from TM tissue dissected out of human donor anterior segment, corneal rims or whole globes. Though these primary cells are utilized for *in vitro* studies in early passages (*∼*2-6), there is a growing recognition among the community that cellular identity is heterogenous, may change with time, and variable from donor-to-donor. Furthermore, there is a considerable interest in the concept of mechanical memory of cells that may help define cellular identity and phenotypic characterizations.

In this study, primary hTM cells were isolated and cultured from non-glaucomatous and glaucomatous donors following tissue dissection, validated through cobblestone morphologic appearance and dexamethasone induced myocillin, and compared with freshly dissected human TM tissue via single cell transcriptomics. Several TM cell specific marker genes were identified, e.g. Chitinase 3 Like 1 (*CHI3L1*), matrix gla protein (*MGP*), and myocilin *MYOC* albeit within different clusters. Prior comprehensive transcriptomic analysis demonstrates the aforementioned genes to be present in the TM and other ocular tissues^15–18^ affirming that the cells characterized are indeed from the appropriate tissue isolated (see Figure 1 for workflow). When the transcriptome of these primary hTM cells were superimposed with those of TM tissue, a striking divergence in cell composition was observed. Specifically, we observe dramatically reduced cell heterogeneity and changes of transcriptomics profiles in the *in vitro* culture compared to *in vivo* tissue.

**Figure 1.**
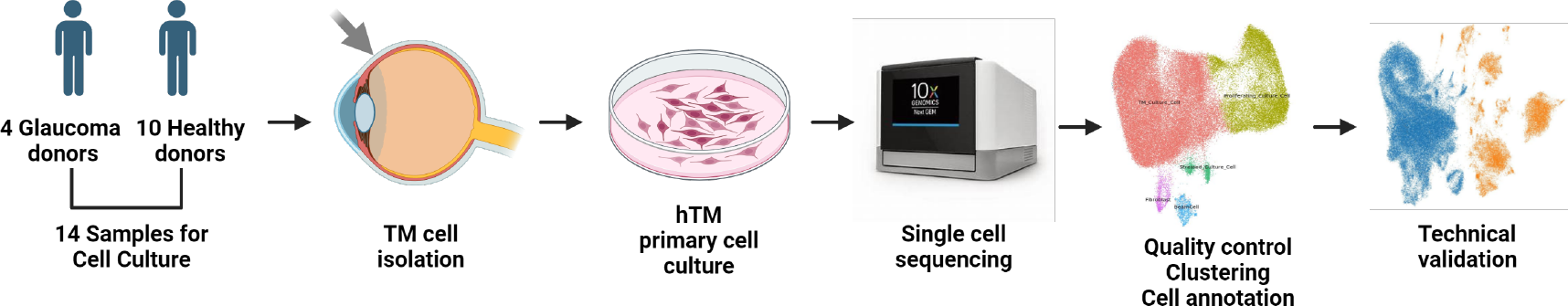
Schematic of experimental workflow

## Methods

### Isolation of donor primary hTM cells

Primary hTM cells were isolated from human donor corneal rims deemed unsuitable for transplantation, and characterized as described previously^19,20^. All donor tissues for primary cell culture were procured from Saving Sight eye bank (Kansas city, MO). All primary cell culture donor cells (Table 1,Table 2) used in this study were isolated (between 2012 and 2022), validated, stored in liquid nitrogen, or used in prior studies from our lab^14,21–24^. Frozen cells were thawed with 15 mL pre-warmed media (37^*?*^C, DMEM:F12=1:1, with 20% FBS, 1% Penn/Step/fungizone) and centrifuged at 300g, 4^*?*^C for 5 minutes. Cells were washed with media again and suspended in 0.04% BFA. Cell viability was checked with trypan blue.

**Table 1.** Donor information and sample IDs for primary cell culture samples.

| Donor_ID | Sample_ID | Sample Type | Disease | Side | Passage |
| --- | --- | --- | --- | --- | --- |
| GTM_1672 | 10x3V31_GTM_1672_P3 | primary cell culture | glaucoma | unknown | P3 |
| GTM_2956 | 10x3V31_GTM_2956 | primary cell culture | glaucoma | OS | P1 |
| GTM_7445 | 10x3V31_GTM_7445 | primary cell culture | glaucoma | unknown | P4 |
| hTM_0114 | 10x3V31_hTM_0114_P1 | primary cell culture | healthy | OS | P1/P2 |
| hTM_11701 | 10x3v31_hTM_11701_P2 | primary cell culture | healthy | OS | P2 |
| hTM_11703 | 10x3v31_hTM_11703_P2 | primary cell culture | healthy | OS | P2 |
| hTM_2180 | 10x3v31_hTM_2180 | primary cell culture | healthy | OD | P1 |
| hTM_659 | 10x3V31_hTM_659_P1_DCR | primary cell culture | healthy | unknown | P1/P3 |
| hTM_74M | 10x3V31_hTM_74M_P2 | primary cell culture | healthy | OD | P2 |
| hTM_7987 | 10x3V31_hTM_7987_P2 | primary cell culture | healthy | unknown | P1/P2 |
| hTM_9355 | 10x3v31_hTM_9355_P1 | primary cell culture | healthy | OS | P1 |
| hTM_9632 | 10x3v31_hTM_9632_P2 | primary cell culture | healthy | unknown | P2 |
| hTM_9691 | 10x3v31_hTM_9691_P1 | primary cell culture | healthy | unknown | P1 |
| OHSU_GTM_2019_0461 | 10x3V31_OHSU_GTM_2019_0461 | primary cell culture | glaucoma | unknown | P3 |
| 22_0500_TM | 22_0500_TM | Post-mortem tissue | healthy | unknown | NA |
| 22_0688_TM | 22_0688_TM | Post-mortem tissue | healthy | unknown | NA |
| 22_0769_TM | 22_0769_TM | Post-mortem tissue | healthy | unknown | NA |

**Table 2.**
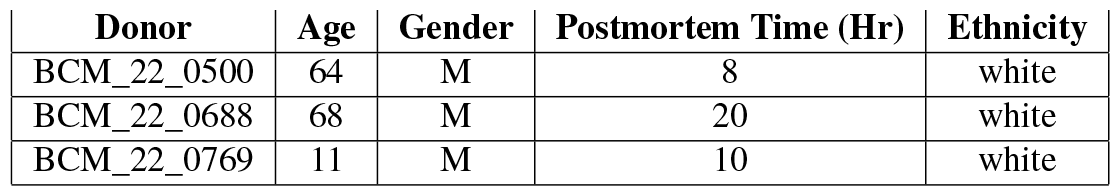
Donor information for post-mortem TM samples.

| Donor | Age | Gender | Postmortem Time (Hr) | Ethnicity |
| --- | --- | --- | --- | --- |
| BCM_22_0500 | 64 | M | 8 | white |
| BCM_22_0688 | 68 | M | 20 | white |
| BCM_22_0769 | 11 | M | 10 | white |

### Donor Tissues

TM tissue for primary cell culture was dissected from donor anterior segments/whole globes obtained from Lions Eye Bank (Baylor College of Medicine, Houston, TX) within 4-6 hours post-mortem. Dissection was performed in accordance with consensus guidelines^19^.

### Single-cell RNA sequencing

Resuspended single cells were loaded on a 10X Chromium controller for obtaining single cell Gel Beads-In-Emulsions. scRNA-seq libraries were generated using 10X Chromium Single Cell 3’ reagent kits v3.1 (10X Genomics) following the manufacturer’s recommendations (https://www.10xgenomics.com). Sequencing was performed on Illumina Novaseq 6000 (http://www.illumina.com) at the Single Cell Genomics Core at Baylor College of Medicine.

### Meta-analysis of scRNA-seq datasets

#### Preprocessing of scRNA-seq Datasets

Raw sequencing reads were processed using the CellRanger v6.1.2 (10X Genomics) pipeline, against the hg38 reference genome (https://cf.10xgenomics.com/supp/cell-exp/refdata-gex-GRCh38-2020-A.tar.gz). Then, quality control was performed separately for each sample through a quality control pipeline (https://github.com/lijinbio/cellqc)^25,26^. Real cells were filtered by dropkick v1.2.8^27^ and ambient RNA was removed using SoupX v1.6.2^28^. Doublets were detected and removed using DoubletFinder^29^. After merging all samples, cells were further filtered manually (features > 300, transcript (UMI) counts > 500, mitochondrial percentage < 5%)(Table 3).

**Table 3.** Cell count after each QC step. *Common cells between Dropkick and Cellranger outputs were used for the next step of QC.

| SampleID | Dropkick | SoupX | Common | DoubletRemoval | Final Filtering |
| --- | --- | --- | --- | --- | --- |
| 10x3V31_GTM_1672_P3 | 10380 | 11699 | 8791 | 8197 | 6910 |
| 10x3V31_GTM_2956 | 6400 | 8611 | 5007 | 4814 | 3257 |
| 10x3V31_GTM_7445 | 10280 | 9139 | 8286 | 7758 | 6943 |
| 10x3V31_hTM_0114_P1 | 8473 | 9162 | 6505 | 6179 | 5682 |
| 10x3v31_hTM_11701_P2 | 8285 | 11929 | 7391 | 6971 | 5025 |
| 10x3v31_hTM_11703_P2 | 7878 | 18391 | 7360 | 6943 | 6012 |
| 10x3v31_hTM_2180 | 16985 | 19081 | 16615 | 14491 | 13340 |
| 10x3V31_hTM_659_P1_DCR | 7312 | 9491 | 7023 | 6644 | 991 |
| 10x3V31_hTM_74M_P2 | 12131 | 12384 | 9591 | 8883 | 7839 |
| 10x3V31_hTM_7987_P2 | 4945 | 7098 | 4101 | 3972 | 3709 |
| 10x3v31_hTM_9355_P1 | 9347 | 11560 | 7951 | 7465 | 6071 |
| 10x3v31_hTM_9632_P2 | 7113 | 8576 | 6563 | 6232 | 3547 |
| 10x3v31_hTM_9691_P1 | 7328 | 8545 | 6687 | 6343 | 4678 |
| 10x3V31_OHSU_GTM_2019_0461 | 12754 | 13404 | 9827 | 9084 | 8539 |

#### Data Integration and Clustering

The raw counts from each sample were merged, and the scVI model (n_layers=2, n_latent=30) from scvi-tools v0.19.5^30^ was used to infer a latent space of 30 dimensions from the raw counts of the top 2000 highly variable genes (HVGs) calculated by scanpy v1.9.1^31^. Sample ID was provided as a batch covariate for the scVI model. The latent space was further reduced using UMAP (min_dist=0.3), and leiden clustering (resolution=0.5) was performed on the scVI latent space, using a k-nearest-neighbor graph (neighbors=20).

#### Cell Clustering and Cell Type Annotation

Cell cycle effects were removed using Seurat^32^. The clustering resolution was validated using sccaf^33^. Initial cell type annotation was performed using scPred^34^ with a reference dataset generated from in vivo tissue samples from our lab. Clusters were re-annotated using known cell type marker genes and cell types were assigned to each cluster using scanpy^31^ with specific markers from previous publications^15,16^. The detailed markers for each cell type are listed in Table 4. Cell proportion plots were generated using dittoseq^35^.

**Table 4.** Highly expressed genes of individual cell type from previous publications.

| Cell Type | Highly-Expressed Genes |  |
| --- | --- | --- |
| Macrophage | C10B, MSR1, MARCO |  |
| T/NK | NKG7, KLRB1, CD3D |  |
| Epithelium | KRT5, SFN, AOP5 |  |
| Melanocyte | MLANA, PMEL, TYRP1 |  |
| Schwann-like | L1CAM, NGFR, SOX2 |  |
| Myelinating Schwann | NGFR, SOX2, PLP1, MPZ |  |
| TM1 (fibroblast-like) | DCN, PDGFRA, TAGLN |  |
| TM2 (myofibroblast-like) | DCN*, PDGFRA, TAGLN, ACTA2, RGS5 |  |
| Smooth Muscle | TAGLN, ACTA2, DES, MYH11, RGS5 |  |
| Pericyte | DCN, TAGLN, ACTA2, RGS5, PDGFRB, FLT1 |  |
| Vascular Endo | RGS5, VWF, FLT1, DLL4, KDR, PECAM1 |  |
| Lymphatic-like Endo | KDR, PECAM1, MMRN1, FLT4, PROX1 | Regneron |
| K-Epi | PAX6 | Sanes |
| Schwalbe Line | CA3, MGARP, ADRB2, SLC2A1, IGFBP2 |  |
| Fibroblast | COL14A1, ADH1B, FBLN2, COL1A2, COL6A1, TNXB, TIMP2, FBLN1, DCN, PCOLCE |  |
| CC | PLVAP, FLT4, PROX1, ACKR1, AQP1, SELE, MMRN1, PECAM1, VWF |  |
| Vascular Endothelium | ALPL, SLC2A1, CLDN5 |  |
| SC | TFF3, PLAT, FN1, POSTN, CCL21, CDH5 |  |
| JCT | NGPTL7, PDPN, CEMIP, MYOC, CYTL1, CHI3L1, NEB, RSPO4, FMOD, NELL2 |  |
| BeamA | BMP5, MGP, RARRES1, FABP4 |  |
| BeamB | TMEFF2, PPP1R1B, BAMBI |  |
| CM | CHRM3, DES, CNN1, MYH11, MYLK, ATP2A1 |  |
| Pericyte | NDUFA4L2, FABP4 |  |
| Neuron | CHRNA3, CALB2, UCHL1, SCG2, GAP43 |  |
| Melanocyte | MLANA, PMEL, MITF, TRPM1, TYR |  |
| Myelinating Schwann | MBP, MPZ, PLP1 |  |
| Non-Myelinating Schwann | LGI4, CDH19 |  |
| BCell | CD27, CD79A, IGHM, MZB1 |  |
| Macrophage | LYVE1, CD68, CXCL8, IL1B, TREM2 |  |
| Mast Cell | RGS13, KIT, CD2 |  |
| NK/T | CD3D, IL7R, TRAC, NGK7 |  |

#### Differentially Expressed Gene Analysis

To identify genes that are differentially expressed between cell types, genes specifically expressed in each cluster were identified and the top 5 genes expressed in each cluster were ranked using Seurat^32^.

#### Data Integration with in vivo Dataset

Cell type of three samples of *in vivo* hTM cells was determined with ScPred individually and then an integrated object of only *in vivo* data was compared with the same known cell type marker genes to determine validity of the *in vivo* samples. Data integration with the tissue culture data was then performed following the same parameters and protocol as above. Then, the combined tissue + culture object was once again compared with the known cell type marker genes and a disparity in gene expression was observed.

## Data Records

Raw reads of all samples and processed data files including integrated data were deposited in the Gene Expression Omnibus (GEO, https://ncbi.nlm.nih.gov/geo) of the National Center for Biotechnology Information (NCBI) as FASTQ files with accession number GSE263230.

## Technical Validation

Prior to sequencing experiments, hTM cells were simultaneously validated by documenting *Myoc* mRNA expression in response to 100 nM dexamethasone treatment for 3 days (Figure 2). Quality control was performed on each sample independently with an average of 6000 cells profiled, yielding a total of about 82k cells (Table 3). There are an average of 2300 gene counts and 8400 UMI counts over the 14 samples (Figure 3(a)). After integration, we affirmed that batch effects were not significant (Figure 3(b)).

**Figure 2.**
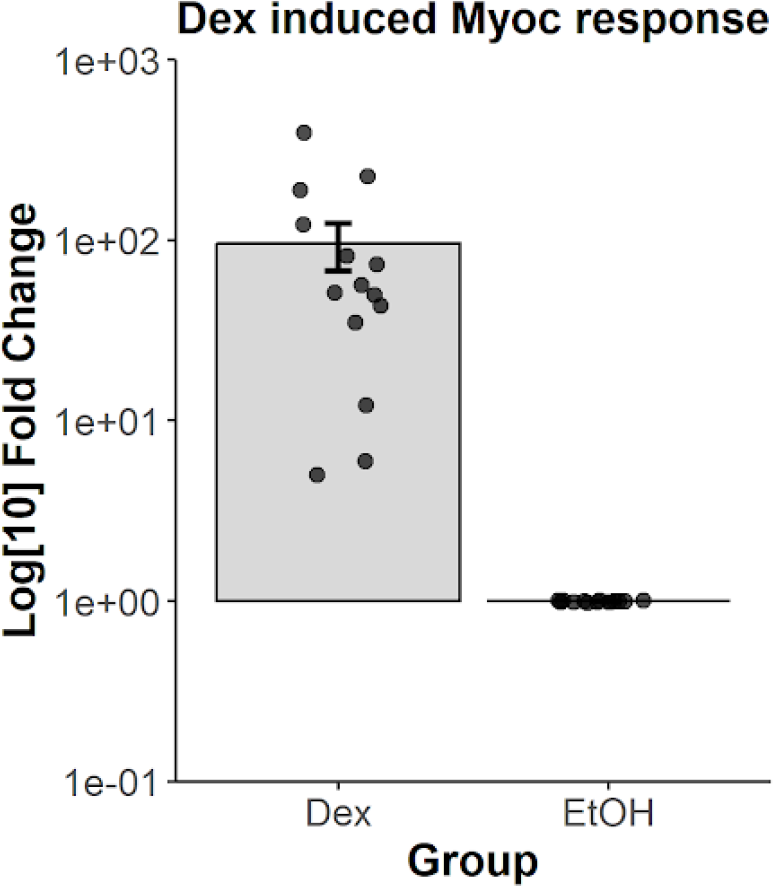
Primary hTM cells used in study demonstrated elevated *MYOC* expression in response to 100 nM dexamethasone treatment for 3 days. Data are from n=14 donors represented as a bar graph, mean *±* standard error in mean. ***p<0.0041, t-test.

**Figure 3.**
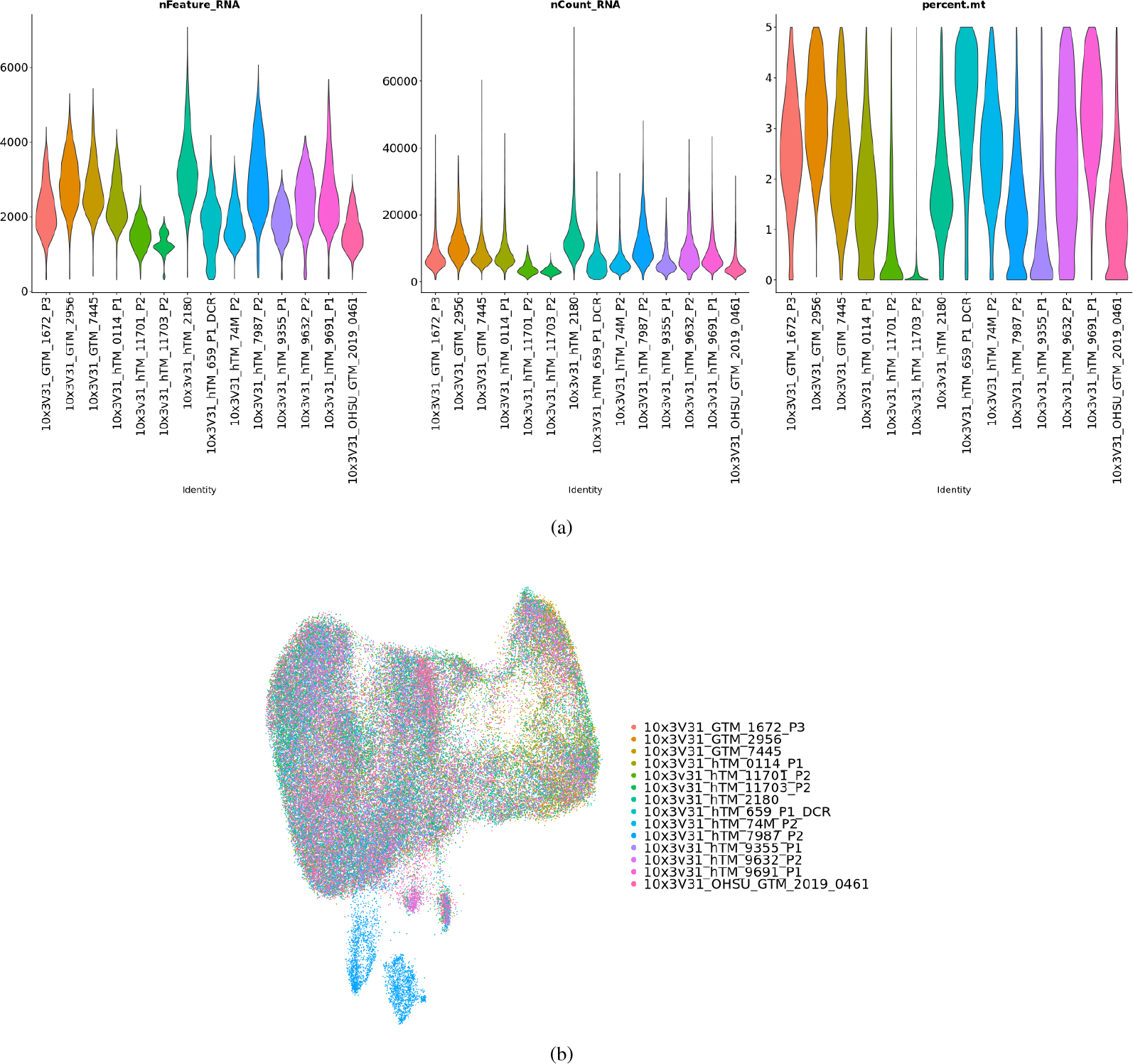
Quality Validation and Characterization of Cells. (**a**) Violin plots showing number of features, number of counts, and mitrochondria percentage by sample. (**b**) The distribution of cells by sample.

By performing clustering analysis of all the cells, a total of 5 clusters were obtained. Of the 5 clusters identified, 3 clusters (TM_Culture_Cell, Proliferating_Culture_Cell, and Stressed_Culture_Cell) are shared by all donors (Figure 4(a)). The other two clusters are only present in the hTM_7987 sample (Figure 4(b)). This suggests that the population of hTM cells in culture is non-uniform and exhibits some heterogeneity. Several TM cell specific marker genes were identified; e.g. CHI3L1, MGP, and MYOC within different clusters. While MYOC was primarily expressed in cluster 0, CHI3L1 and MGP were abundant in cluster 3. When overlaid with cell-specific markers previously reported, PDPN, CEMIP and CHI3L1 were abundant in cluster 3, while MYOC remained enriched in cluster 0. Matrix proteins, collagens 1/6, FBLN, FN, POSTN, and DCN were enriched in clusters 3 and 4, while PCOLCE was enriched in clusters 0 and 1 (Figure 4(c)). Prior comprehensive transcriptomic analysis demonstrates the aforementioned genes to be present in the TM and other ocular tissues^15–18^ affirming that the cells characterized are indeed from the appropriate tissue isolated.

**Figure 4.**
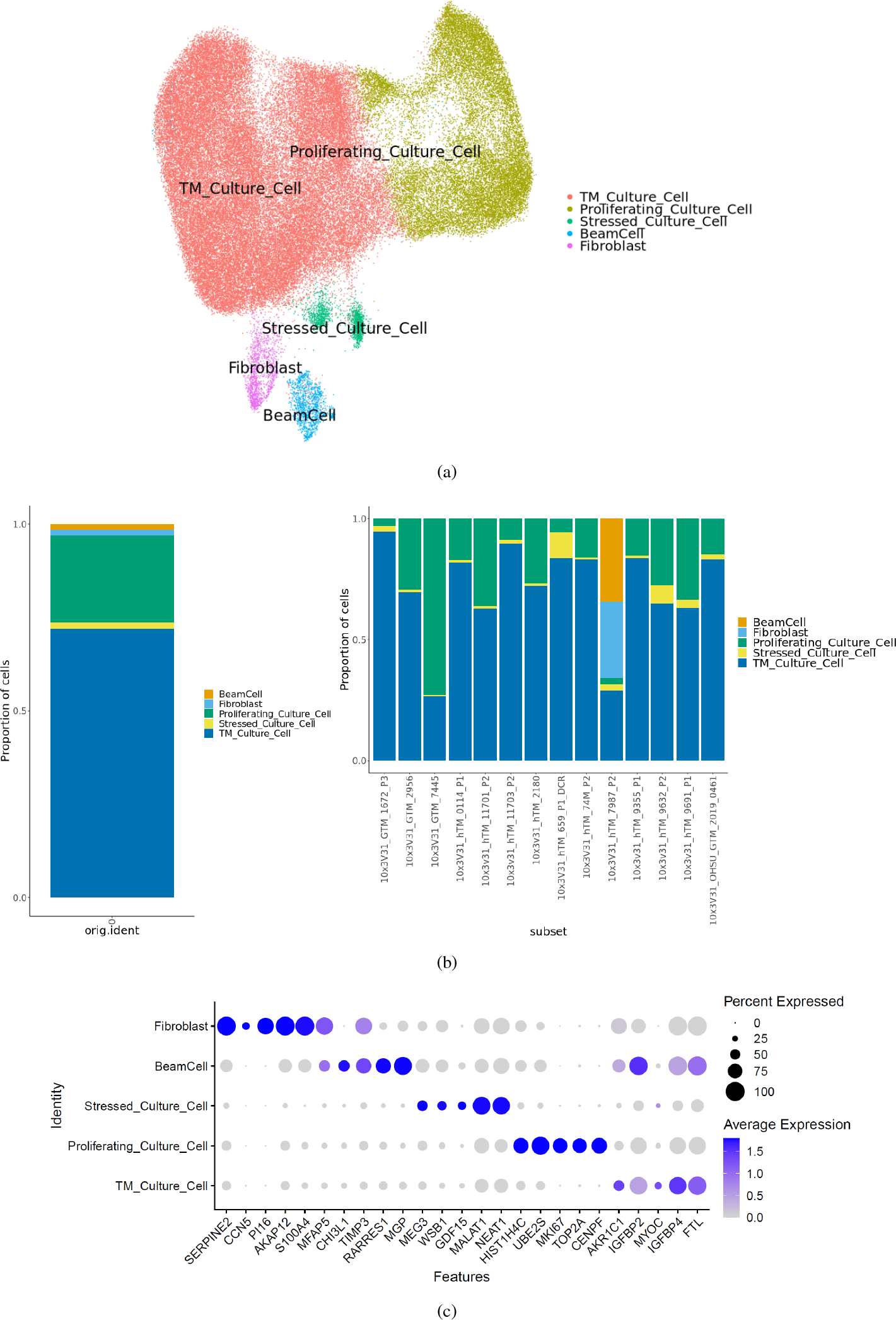
Quality Validation and Characterization of Cells. (**a**) Umap of integrated object with cell type annotations. (**b**) Cell proportion bar plot. (**c**) Dotplot with top 5 highly expressed genes

With the tissue data, we used the same QC pipeline with the same parameters for prepossessing. In the tissue samples alone, there are an average of 8100 cells profiled, yielding a total of about 24k cells with an average of 2300 gene counts and 6400 UMI counts. We assigned cell types using scPred to yield 9 defined cell types with a few unassigned cells (Figure 5(a)). We validated the clustering and cell type assignment with previously identified marker genes^15–18^ and confirmed that the data were good for further analysis.

**Figure 5.**
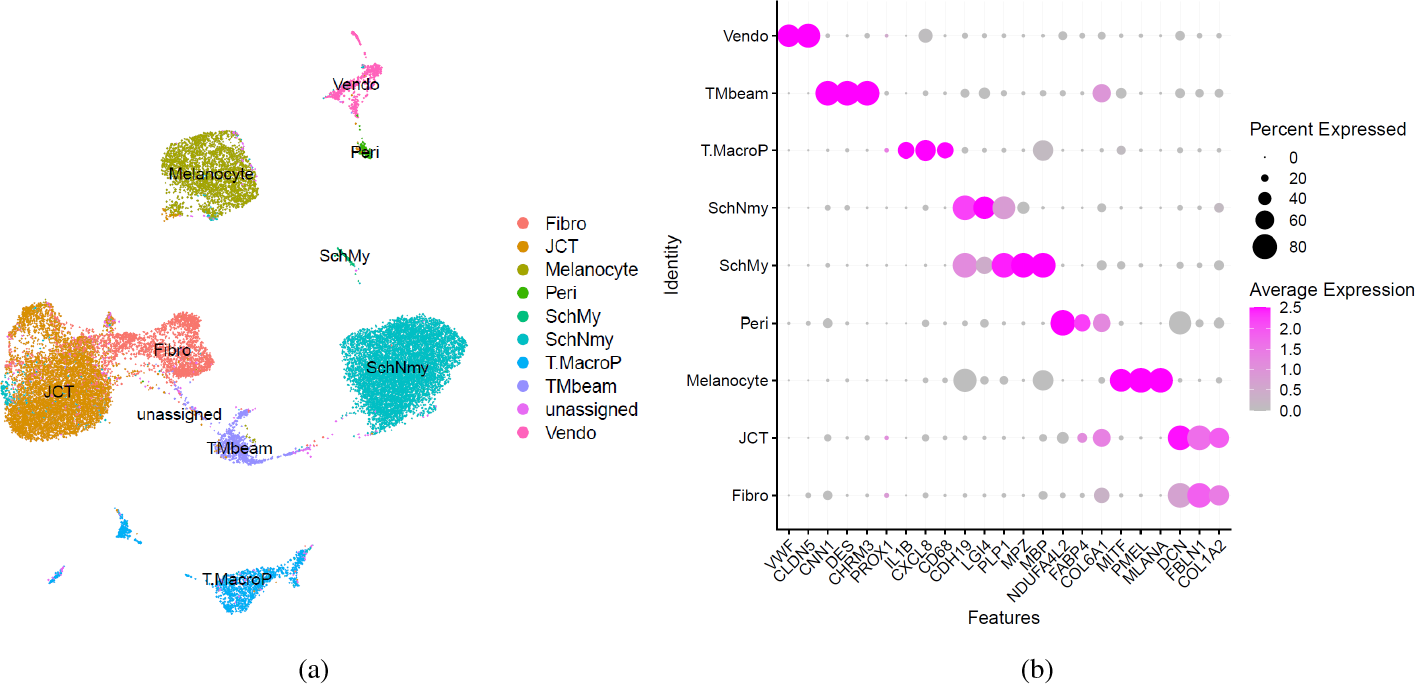
Single cell transcriptome of TM tissues for quality Validation. (**a**) UMAP of *in vivo* hTM cells with cell type annotation by Seurat. (**b**) Dot Plot showing marker gene expression by cluster

## Usage Notes

Our dataset will be useful for a variety of studies pertaining to understanding the identity of primary hTM cells, including studies to translate pre-clinical *in vitro* models to *in vivo* models mimicking disease, relevant biophysical and biochemical cues mimicking the native cellular microenvironment, and choice and type of *in vitro* models utilized for investigations. Here we provide a comparison between our dataset and tissue data generated by our lab as a usage example:

When the transcriptome of these primary hTM cells were superimposed with those of TM tissue, a striking divergence in cell identity was observed. Specifically, reduced cell heterogeneity and dramatic changes of transcriptomics profiles in the *in vitro* culture compared to *in vivo* tissue (Figure 6). It is important to note that the primary hTM cells characterized in this study were previously frozen down in liquid N2 and are from early passages (up to 2). It is unclear why and at what stage these putative markers may be lost in culture. We are also uncertain, if at any time in culture, these cells may have de-differentiated from a select population of proliferating cells since heterogeneity while observed *in vitro* is lesser compared with those in the tissue.

**Figure 6.**
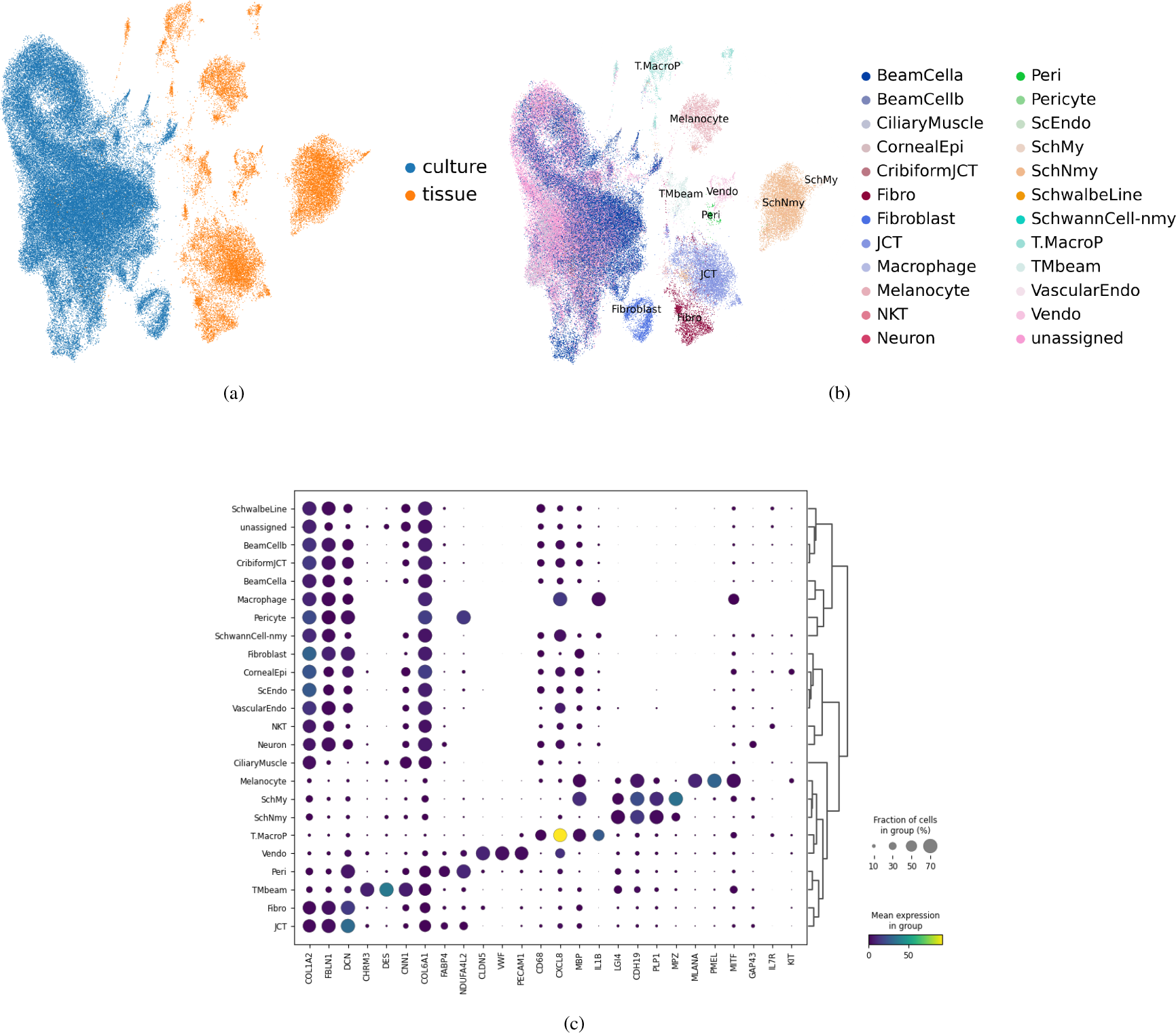
Data integration for comparison of *in vivo* and *in vitro* data. (**a**) UMAP by sample type with separation of clusters between primary cell culture and post-mortem tissue. (**b**) UMAP by cell type with separation of clusters between primary cell culture and post-mortem tissue. (**c**) Dot Plot showing marker gene expression by cluster between both sample types

Our group and others have consistently reported that biophysical and biochemical cues from the cell culture microenvironment (topography, stiffness, ECM coating, stretch, 2D vs 3D) have profound impact on transcriptomic, proteomic, signaling pathways, and response to drugs in vitro^36–55^. Thus it is feasible to infer that while mechanical cues may drive hTM cell function as a function of substrate properties, the initial culture conditions in which these cells were first isolated and expanded may have also profound impact on selection of cells for propagation, proliferation, and (de-)differentiation. That mechanical memory and plasticity exists in maintaining cell identity through epigenetic regulation has been previously postulated^56–62^. However, whether such a phenomenon exists in primary hTM cells remains to be further determined. Further, emerging evidence suggests chromatin remodeling implicating significant changes to cellular and epigenetic plasticity may significantly alter cell fate with prolonged exposure to rigid culture environments such as tissue culture plastic^61–64^. Since all primary hTM cells are primarily cultured and expanded on rigid plastic substrates, it is only natural to infer that the process of divergence in cell identity as observed in this study likely starts immediately after initial isolation. A systematic study is critically needed to confirm this, and this may subsequently allow for the development of appropriate culture conditions and microenvironments which better maintain primary TM cell identity.

All raw RNA sequencing data are stored in FASTQ files, and processed .h5ad files are also available for use.

## Code availability

The source code, including code to generate all figures, has been uploaded to GitHub: https://github.com/RCHENLAB/TM_culture_manuscript.

## Acknowledgements

This project was funded by NIH/NEI R01EY022356 (R.C.), R01EY018571 (R.C.), S10OD032189 (R.C.), R01EY026048-01A1 (V.R.), Chan Zuckerberg Initiative (CZI) award CZF2019-002425, RRF to R.C. We would also like to thank the Lions Eye Bank (TX), Lions Vision-Gift (OR), and SavingSight (MO) for procuring all human donor eyes used in this work. Most importantly, we would like to thank the families of the organ donors without whose consent these experiments would be impossible. Janice A Vranka passed away in 2021 after a battle with cancer; her intellectual contributions to the study are invaluable

## Author contributions statement

A.T., V.R., and R.C conceptualized and designed the study. Y.L. generated the scRNA-seq datasets in this study. All authors wrote, reviewed, and contributed to the manuscript.

